# DNA Compass: a secure, client-side site for navigating personal genetic information

**DOI:** 10.1101/077867

**Authors:** Charles Curnin, Assaf Gordon, Yaniv Erlich

## Abstract

**Motivation:** Millions of individuals have access to raw genomic data using direct-to-consumer companies. The advent of large-scale sequencing projects, such as the Precision Medicine Initiative, will further increase the number of individuals with access to their own genomic information. However, querying genomic data requires a computer terminal – an impediment for the general public.

**Results:** DNA Compass is a website designed to empower the public by enabling simple navigation of personal genomic data. Users can query the status of their genomic variants for over 400 conditions or tens of millions of documented SNPs. DNA Compass presents the relevant genotypes of the user side-by-side with explanatory scientific resources. The genotypes data never leaves the user’s computer, a feature that provides improved security and performance. Nearly 2500 unique users have used our tool, mainly from the general genetic genealogy community, demonstrating its utility.

**Availability:** DNA Compass is freely available on https://compass.dna.land.

## 1 Introduction

We have entered the era of ubiquitous genomic information. Today, approximately three million people worldwide have access to their genome-wide autosomal information via direct to consumer (DTC) genomic companies such as 23andMe, AncestryDNA, and FamilyTreeDNA (Khan and Mittelman, 2013). Recent studies have predicted that by 2025, at least 100 million individuals will have their genomes sequenced (Stephens *et al.*, 2015) and whole genome sequencing will become a routine part of newborn screening (Burn and Flinter, 2013).

Interpretation of this data, however, remains difficult. While recent ethical standards have highlighted the importance of returning results to research participants, most studies are reluctant to provide any interpretation to the participants due to regulatory complications (Jarvik *et al.*, 2014; Evans and Rothschild, 2012). As an alternative, a growing number of studies, including DTC companies or Genes for Good, return raw data to participants. However, the scale of genomic information precludes even simple analysis by people with no bioinformatics skills. VCF files of personalized genomic tests can reach gigabytes of data; although these files are textual and human-readable, they are largely inaccessible to individuals who lack basic command-line skills. We encountered this problem recently with our website DNA.Land (https://dna.land) (Erlich, 2015), which crowdsources genomic datasets directly from people who were tested by DTC companies.

One of the features of the website includes genomic imputation of the user’s DTC file to report 39 million variants. We anticipated that this unique feature would be well-received by the 30,000 participants of DNA.Land. However, we found that most participants who downloaded the data expressed a high level of frustration after repeatedly crashing their Excel spreadsheet or Microsoft word processor when attempting to open the imputed VCF file.

Here, we present DNA Compass (https://compass.dna.land), a free website that enables the navigation of personal genomic information without command-line knowledge. Our website aims to empower the growing number of individuals who have access to their genomic datasets but are uninterested in pursuing a bioinformatics or computer science degree just to search for a particular SNP in their raw data. To test the website, we announced its launch on the Facebook pages of several genetic genealogy groups. We had nearly 2500 unique users within the first two days of operation, demonstrating the wide interest in such a tool. Most users were able to process their data easily and, overall, we received very positive feedback.

## 2 User experience

Users begin interacting with DNA Compass by uploading two files: a compressed VCF file and a corresponding Tabix index file (Danecek *et al.*, 2011). We selected these file formats since they are compatible with the files reported to DTC participants on DNA.Land and due to their wide popularity in whole genome sequencing experiments.

Next, DNA Compass quickly processes the genomic information, validates the format, and presents the user with a report through which they can navigate their genetic information. The user only needs to enter a particular condition (e.g. height or skin pigmentation) from more than 400 conditions. To assist the user to find the condition of interest quickly, the website completes the text as the user types and also offers the full list of conditions. Next, DNA Compass presents a table that lists the dbSNP rsID, the user’s genotype, and chromosome for significant (p<10^−15^) SNPs that are associated with the condition according to NCBI’s Phenotype-Genotype Integrator (Ramos *et al.*, 2014). Importantly, the website does not directly interpret genomic information. Instead, for each SNP, it collates five publicly available resources, namely SNPedia (Cariaso and Lennon, 2012), PubMed, dbSNP (Sherry *et al.*, 2001), GWAS Central (Beck *et al.*, 2014), and Google. With the exception of Google, clicking on any of these resources opens an iframe window that presents the selected website side-by-side with the user’s own genetic information (Figure 1). Crucially, we do not send any private information to any of these websites. This model allows the user to access relevant scientific knowledge with maximum efficiency. If the user chooses the Google search option, a new tab is opened in the browser where the search term is the SNP’s rsID. In addition to navigating SNPs based on traits, the website also allows users to enter particular SNPs (e.g. rs10037512).

**Figure 1:**
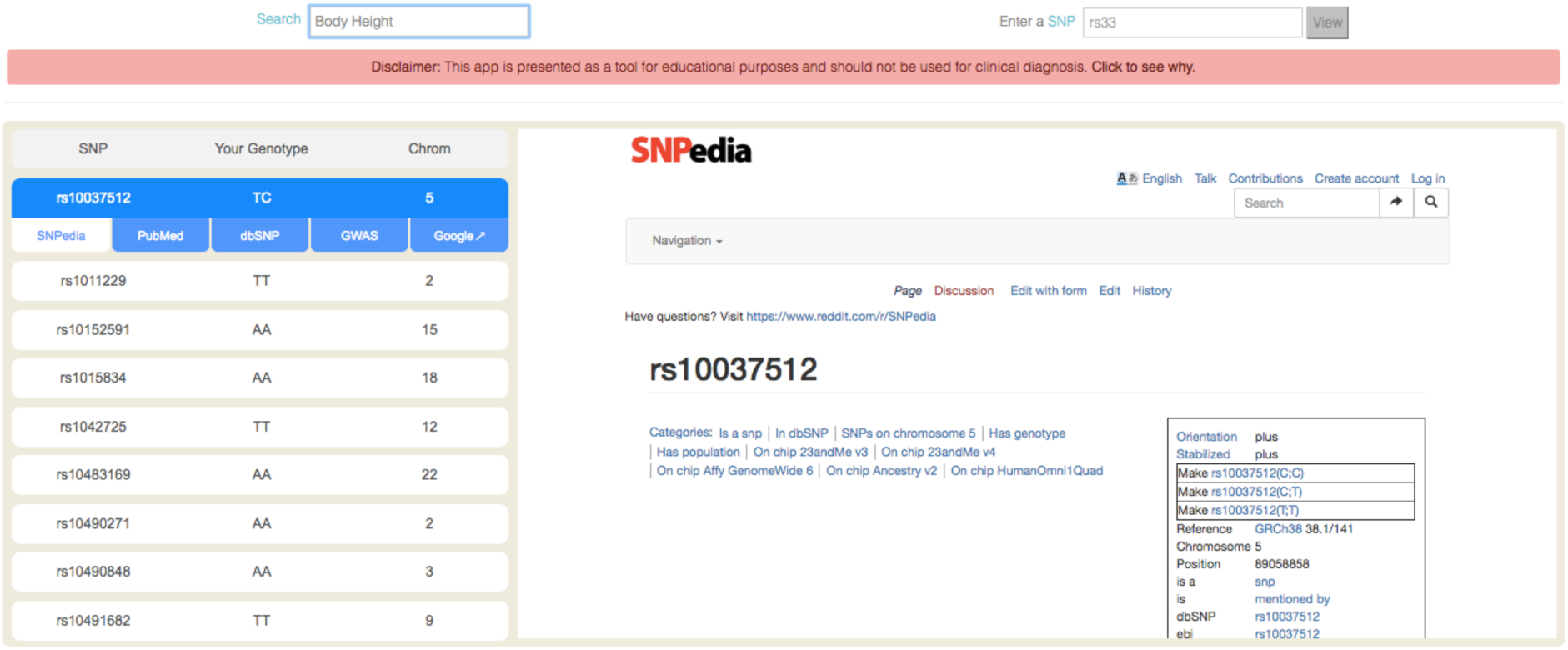
DNA Compass report for some of the height-associated genotypes of a user (left) along with information from SNPedia (right). By clicking on each SNP in the list (left), the user can select relevant information related to the SNP from SNPedia, PubMed, dbSNP, GWAS Central, or Google. The search field (top left) supports over 400 conditions. The SNP field (top right) allows the query of tens of millions of variants. The data presented belongs to the senior author of this manuscript.

There are several options for user support. First, an example of VCF-Tabix pair is available for those who want to test the website. In addition, we include an extensive FAQ section and guide. Finally, the website is dynamic and provides quick feedback for the user’s actions.

## 3 Architecture

DNA Compass operates almost exclusively on the client-side. The sole role of the server is to provide static webpages and information regarding SNP positions and associations. Importantly, the client never transmits information from the user’s VCF and Tabix files. This has two advantages. First, it obviates the need for transferring massive VCF files across the web, which means a quick response to user queries. Tests with a standard Macbook Air laptop show that it takes a few seconds on either Chrome or Firefox to return the results of a query. Second, by restricting the processing of data to the client side, we mitigate genetic privacy issues related to the management and storage of personal genomic data (Erlich and Narayanan, 2014).

The server-side uses DreamFactory and SQLite3 to store the coordinates (chromosome and GRCh37/hg19-compliant position) for each SNP rsID according to dbSNP141. The client-side functionality employs jQuery, Bootstrap, and jsf-local-aerial. To facilitate future developments, we also released the entire source code on GitHub (https://github.com/TeamErlich/dna-land-compass) under the BSD license.

## 4 Conclusion

With the advent of large-scale public efforts to sequence a massive number of individuals and the increasing integration of genomics in medicine and self-curiosity, we envision a growing demand among the general public for technological solutions that will allow them navigate their raw data. DNA Compass aims to increase genetic literacy and empower research participants to understand their personal genomic data.

## Acknowledgements

We thank that DNA Compass users who tested the website and helped us to further develop the website and Dina Zielinski for helpful comments.

